# ThermoClock:A Novel Automated Temperature Regulation Device that Can Model Circadian Entrainment and Disruption in 2D and 3D in Vitro Models

**DOI:** 10.64898/2026.07.30.741626

**Authors:** Kaiyin Kelly Zhang, Cathryn A. Cutia, Chanelle A. Moise, Bhavna Kalyanaraman, Kaitlyn Chhe, Jiexin Wang, Michelle E. Farkas, Ilia N. Karatsoreos, Mary E. Harrington, Meghan E. Huber, Cathal J. Kearney

## Abstract

Circadian rhythms are critical for maintaining homeostasis and regulating physiological functions, and consequentially impact disease progression; yet, they remain largely overlooked in in vitro models used in preclinical research. One major barrier to rigorously testing the role of circadian rhythms in these models is the lack of accessible tools that seamlessly integrate into standard culture setups and are capable of sustainably delivering time cues to cells and tissues in long term experiments. Here, we present the ThermoClock, a low-cost, Arduino-based automated temperature control system capable of delivering independent temperature programs to multiple cultures simultaneously. Using circadian reporter U2OS cell lines (Bmal1:Luc and Per2:Luc), we demonstrated that ThermoClock-driven temperature cycles (36°C/38.5°C, 12h:12h) produced significantly higher amplitude entrainment than a programmable incubator delivering identical temperature trajectories, suggesting that the ramp time to setpoint is a critical determinant of entrainment strength. We further applied ThermoClock to skin explants from keratinocyte-specific Dbp:Luc reporter mice, showing that circadian temperature cycles (T24: 12h:12h and T25: 12.5h:12.5h) extended synchronized circadian rhythms ex vivo, while a shortened T-cycle (T20: 10h:10h) induced rhythm disruptions. We also observed reduced cell migration in T20 temperature-entrained explants wounded ex vivo, closely recapitulating attenuated wound healing observed in T20 light-cycle-disrupted mice in vivo. Finally, we show that wounding can act as a phase-resetting cue but its efficacy depends on pre-injury entrainment state, with circadian entrained tissues (T25) resisting reset, while disrupted (T20) and unentrained tissues showed resetting sensitivity. These findings establish ThermoClock as a versatile platform for incorporating circadian regulation and, for the first time, disruption into 2D and 3D in vitro systems and demonstrate that peripheral clock disruption and its functional consequences can be modeled ex vivo.

## Introduction

Circadian rhythms are driven by an endogenous biological clock that aligns behavior and physiology to an approximately 24-hour cycle. Understanding how circadian variations shape physiology, and how chronic disruption to circadian rhythms impact cellular functions carries significant clinical implications [1], [2]. Time of day is shown to influence outcomes in disease progression [3], [4] , as well as surgery outcomes and therapeutic efficacies [5]. Adding to that, the internal circadian clock that drives these rhythms is becoming increasingly vulnerable to chronic disruption due to modern lifestyles—artificial light exposure at night, irregular mealtimes and shiftwork—exposing individuals to conflicting circadian time signals. Chronic disruption of the clock elevates risk of various major diseases, including obesity, type 2 diabetes, and cardiovascular diseases[6], [7], [8], [9]. Beyond systemic disease, temporal control is equally critical in tissue repair and regeneration. Circadian rhythms govern key aspects of the wound healing responses, with healing rate shown to be dependent on the time of injury onset and attenuated under disruptive rhythms[10], [11].

Even with this demonstrated impact, circadian factors are typically overlooked in therapeutic development and in the clinical translation process, but these are critical to consider for multiple reasons. First, therapeutics are rarely evaluated in circadian disrupted models, though patients of many diseases experience chronic circadian disruption[12], [13]. Second, many drugs show different effects when administrated at different times of day[14]. Third, pre-clinical research procedures are often conducted at convenient hours during natural daytime (light-on hours).

These results might not be translatable for diseases that occur during human’s active phase as nocturnal animals (e.g., rodents used in preclinical research) are on their rest phase during the procedure [8], [15], [16]. Given the critical role of circadian rhythms in physiological processes, it is important to include them in all phases of biological studies, including early stage 2D and 3D in vitro studies. However, the majority of isolated cells and tissues in culture lose cell-to-cell circadian rhythm synchrony rapidly in the absence of coordinated time signals, creating a key discrepancy from their in vivo counterparts[17], [18], [19]. As new approach methodologies (NAMs) such as organ-on-a-chip and other microphysiological systems are built on the same culture principles, the physiological relevance and translational impact of these promising technologies remain fundamentally constrained by the lack of synchronized circadian clocks[20], [21]. Taking the state of organismal, tissue and cellular circadian clocks into consideration in studying physiology, evaluating biomaterials, and developing therapeutics is becoming a necessity to improve translational efficacy.

In in vivo studies, these challenges are more readily addressed: we can house animals in isolated units with controlled light/dark schedules that align the procedure time with the targeted circadian phase. The suprachiasmatic nucleus (SCN) acts as the central pacemaker to rest of the body—it perceives the daily cycles of light, one of the strongest *zeitgebers* (time giving cues), through optical nerves and synchronizes the clocks in peripheral tissues via different signals including temperature and hormonal oscillations[6], [22], [23]. Light cycles of various non-24h periods, “T-Cycles”, have also been used to induce circadian disruptions in behaviors and metabolisms[24], [25]. In vitro systems, which are typically not light sensitive, are instead synchronized by cues mimicking downstream effects of the SCN. In vitro, cells/tissues are synchronized by various inputs such as glucocorticoids [26], serum shock[27], and temperature oscillation[23].

Challenges remain for in vitro circadian entrainment. With both chemical and serum pulses, as peripheral tissues desynchronize within three to four days, it requires repeated stimulation for long-term experiments; but the timing and overlap of repeated pulsing have not been robustly established. Temperature entrainment through multiple cycles between two near-physiological temperatures — such as 36–38.5°C or 32–37°C every 12 hours — has been shown to be an effective and continuous method to entrain circadian rhythms in mammalian explants and cell cultures via activation of heat shock factor (HSF)-mediated transcription [10], [23], [28].

Temperature entrainment demands continuous cycling, yet most commercial incubators are designed to maintain a constant temperature, and they require long ramp times for temperature changes[29]. Both approaches remain labor-intensive for prolonged studies, necessitating the need for advanced equipment, thereby discouraging most researchers from including the circadian component in their studies. Importantly, there is no generally accepted method to model circadian disruption in vitro. This is critical for progress in the mechanistic understanding of circadian disruption, the impact on cellular health, and therapeutic consequences.

In this study, we present the ThermoClock, a novel, low-cost, and automated temperature control system designed for in vitro and ex vivo translational research. Using this system, we successfully entrained circadian rhythms of immortalized reporter cell line and skin explants from novel cell-specific circadian reporter mice. Notably, cell line experiments suggested that the temperature ramp rate influences entrainment outcomes, providing new insights into temperature-based circadian rhythm entrainment. Beyond healthy rhythms, the ThermoClock was also used to induce circadian disruption in vitro for the first time, by using T20 cycles in skin explants and this enabled the isolated study of peripheral clock disturbance and its impact on wound healing. Finally, we investigated the relationship between tissue injury and peripheral clock function via a temperature-entrained ex vivo wound model. This study not only established ThermoClock as a feasible and accessible platform for conducting in vitro circadian entrainment but also demonstrated that circadian variations and disruptions can be present in vitro.

Incorporating both entrained and disrupted circadian oscillations into cell-based models is an important step to improving in vitro physiological models for their use in the therapeutic development pipeline. The ThermoClock is an inexpensive, effective way to achieve this.

## Materials and Methods

### Cell Culture

The circadian reporter cell lines used in this work were established previously, with promotor-driven reporter plasmids of Bmal1 (pABpuro-BluF, Addgene plasmid #46824, deposited by Dr.Steven Brown) and of Per2 (Per2:luc, Addgene plasmid #206875, deposited by Dr.Michelle Farkas). Bmal1 and Per2 are core components of the circadian TTFLs and are expected to oscillate anti-phase with each other[30]. U2OS-Bmal1:luc and U2OS-Per2:luc cells were cultured in high glucose Dulbecco’s modified eagle medium (DMEM, Gibco, Waltham, MA) supplemented with 10% fetal bovine serum (Biowest, Bradenton, FL), 2 mM of L-glutamine (Sigma-Aldrich, St. Louis, MO), 100 U/mL penicillin–streptomycin (Gibco), 1% non-essential amino acids (Cytiva, Marlborough, MA), and 1 mM of sodium pyruvate (Gibco).

### Animals

All animal procedures were approved by the Institutional Animal Care and Use Committee of University of Massachusetts Amherst or Smith College. For bioluminescence recording, 3–4-month-old male and female albino D-domain binding protein(Dbp)KI;tyr conditional circadian reporter mice that were heterozygous for the K14-Cre transgene (Tg(KRT14-cre +) / +, strain#:018964; DBPKI + /DBPKI + ; B6(Cg)-Tyr c-2J) were bred in-house and utilized for this experiment[31], [32]. For all other procedures, 3–4-month-old male and female wildtype C57BL/6 mice were used (Jackson Labs, Bar Harbor, ME). Mice were group housed in 12h:12h light/dark unless otherwise specified, with food and water available ad libitum.

### ThermoClock device design and set up

The ThermoClock’s detailed design, manufacturing instructions and validations results have been previously described[29]. In brief, it is an Arduino-based automated temperature control system, pairing PT100 resistance temperature detectors (RTDs, Adafruit, New York, NY) and electric heating pads (Adafruit) via proportional-integral-derivative (PID) controllers, capable of regulating temperatures for five independent cultures simultaneously. In these studies, we used 35mm cell culture petri dishes (Nunc^®^, Thermo Scientific, MA or Falcon^®^, Corning Inc, Corning, NY) but the ThermoClock is also compatible with multi-well plates of different formats. The ThermoClock was placed inside an incubator set at the lowest temperature of the temperature treatment trajectory (36℃ for this study), and at 5% CO^2^ and default incubator humidity unless stated otherwise.

### Cell culture sample plating and entrainment

Reporter U2OS cells were plated at 100,000 cells per dish in 35mm cell culture petri dishes (Nunc^®^ Thermo Scientific) with 2mL of U2OS culture media. One day after plating, samples in the entrained groups were placed on the ThermoClock, or in a programmable incubator (Binder, Bohemia, NY), to begin entrainment with 3 cycles of 36°C 12h/ 38.5°C 12h.

Dexamethasone-pulsed and no treatment controls were cultured at 37°C throughout the process. Following entrainment, all temperature entrained samples were returned to 37°C. Simultaneously, dexamethasone control samples started incubation with 100nM dexamethasone (Sigma-Aldrich) in fresh media for 2h (Fig.2A). All sample media was then replaced with 2mL of fresh, sterile-filtered recording media for U2OS [33]: powdered DMEM (Sigma-Aldrich), 4mM sodium bicarbonate (Gibco), 5% FBS, 1% HEPES (Sigma-Aldrich), 0.25% penicillin-streptomycin (Gibco), 150μg/mL d-luciferin (ThermoFisher). The dishes were then sealed with vacuum grease (Dow-Corning, Midland, MI) and 40mm glass coverslips (ThermoFisher) for bioluminescence recording.

**Figure 1.**
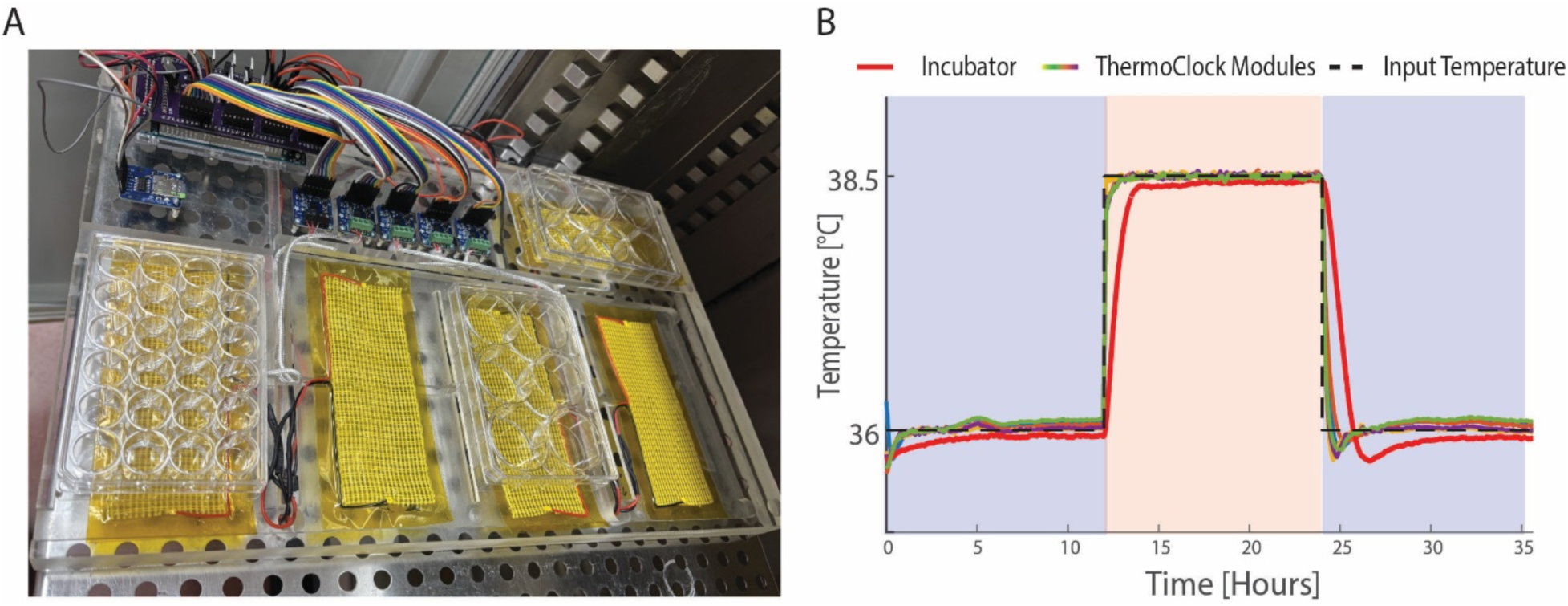
ThermoClock is a cell culture-compatible temperature regulation system with the rapid setpoint change capability. A) Assembled ThermoClock with five modules (pictured). The system is fully functional inside a standard, commercial incubator. B) ThermoClock modules are able to achieve setpoint temperatures, both in cooling and heating, significantly faster than an incubator. Pink region indicates input temperature at 38.5°C, blue region at 36°C.

**Figure 2.**
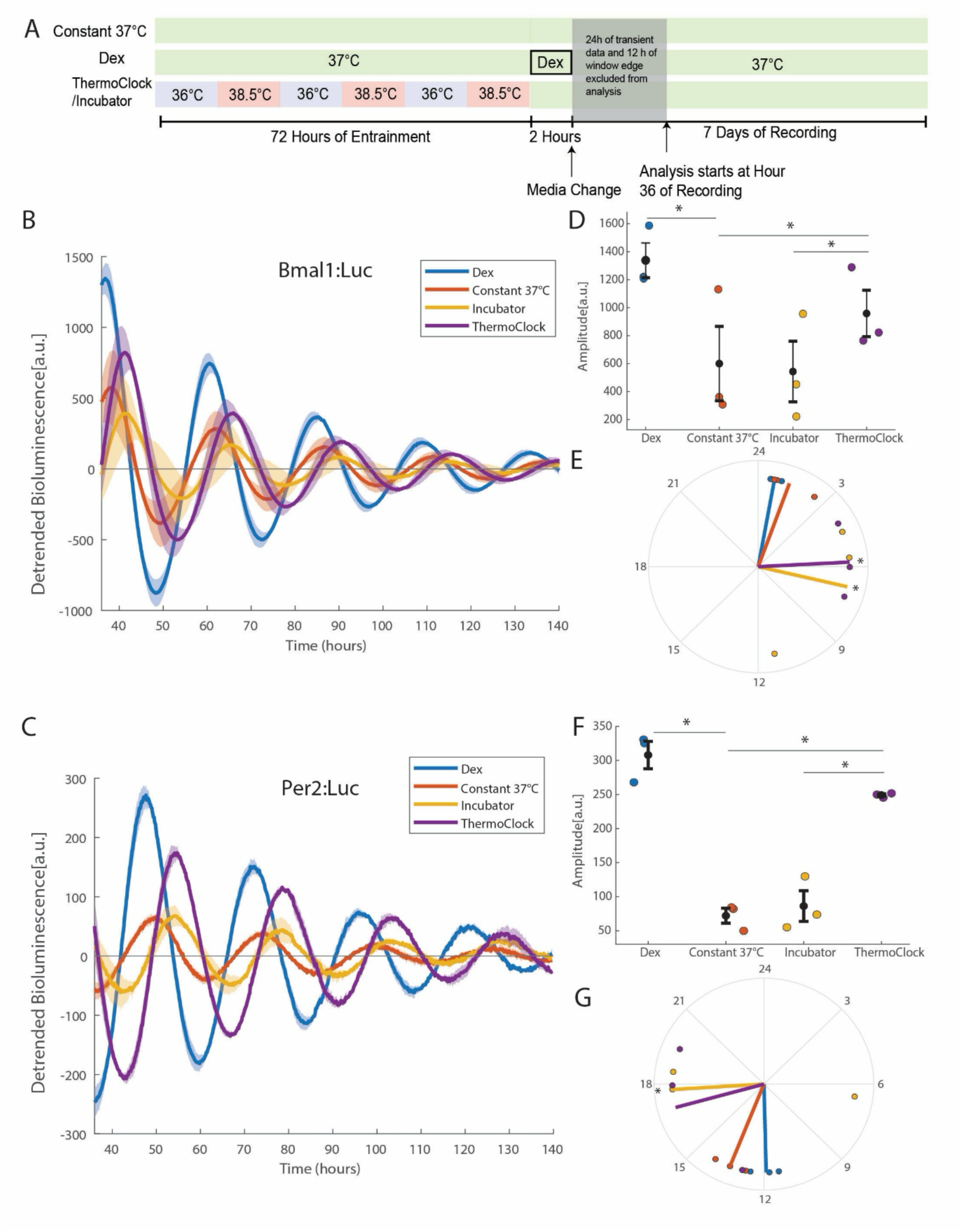
ThermoClock entrained Bmal1:Luc and Per2:Luc U2OS showed robust rhythms. A) Bmal1:Luc and Per2:Luc U2OS cells were entrained by 3 cycles of 12h:12h 36℃(blue)/38.5℃(red) by ThermoClock or incubator. Start of the 2h dexamethasone pulse (positive control) aligned with the end of the temperature entrainment program. All samples were recorded at 37℃(green) for 7 days. B) Averaged detrended Bmal1:Luc U2OS bioluminescence data (N=3; mean±SEM). C) Averaged detrended Per2:Luc U2OS bioluminescence data (N=3; mean±SEM). D) Amplitudes and E) phases of Bmal1:Luc U2OS (fitted from 36h to 116h; N=3, mean±SEM; one-way repeated measures of ANOVA with Sidak’s post hoc test, *p < 0.05) F) Amplitudes and G) phases of Per2:Luc U2OS (fitted from 36h to 116h; N=3, mean±SEM; one-way repeated measures of ANOVA with Sidak‘s post hoc test compared to constant 37, *p < 0.05). *Time in Fig.2B,C is time since the start of recording.

### Skin Explant Preparation and Entrainment

For bioluminescence recording, K14-Cre Dbp:Luc transgene (Tg(KRT14-cre +) / +; DBPKI + /DBPKI +; B6(Cg)-Tyr c-2J) mice were euthanized 2 to 3h after lights on, via isoflurane overdose followed by cervical dislocation. Dorsal skin samples were collected using a 7.0mm biopsy punch (SHARD^®^, AD Surgical, Sunnyvale, CA) after fur had been removed, and sterilized with 70% EtOH. Samples were placed in ice cold Hank’s balanced salt solution (Invitrogen, Carlsbad, CA) and were immediately randomly assigned to different entrainment conditions and plated on a 0.4µm culture insert (Millipore, Billerica, MA) in a sterile 35mm petri dish (Falcon) with 3 ml of bicarbonate-buffered bioluminescence medium[34]: DMEM (Sigma-Aldrich, St. Louis, MO) supplemented with 2% B-27 (Invitrogen), 4 mM L-glutamine (Invitrogen), 10 mM glucose (Sigma), 4.2 mM NaHCO_3_ (Sigma), 10 mM HEPES (Sigma), 25 units/ml penicillin-G sodium (Invitrogen), 34 μM streptomycin sulfate (Invitrogen), and 100 μM beetle luciferin (Promega, Madison, WI). The dishes were then sealed with vacuum grease (Dow-Corning) and a 40mm glass coverslip. The control group was placed directly into the LumiCycle (32 well, Actimetrics, Wilmette, IL) after collection. Entrainment samples were placed on different ThermoClock modules for entrainment with the following temperature cycles: T20 (10h:10h 36°C /38.5°C), T24 Nocturnal (12h:12h 36°C/38.5°C), T24 Diurnal (12h:12h 38.5°C/36°C), and at T25 (12.5h:12.5h 36°C/38.5°C). After three cycles of entrainment, samples were maintained at 37°C until T25 entrainment completed (75h after collection), and then transferred into the LumiCycle (Actimetrics).

### Wounded Skin Explant Preparation

Prior to wounding, explants were collected from wildtype C57BL/6 and entrained identically to the above experiments. Explants entrained by T20, T25 and control samples (no entrainment) were selected for the wounding study. Immediately at the end of T25 entrainment, the explants were wounded with 2mm biopsy punches (Integra, York, NY) in the center. The wound was filled with a fibrin solution (2.5mg/mL fibrinogen and 1U/mL thrombin), which polymerized into a clot-mimicking hydrogel. Wounded skin explants were plated on a culture insert (Millipore, Billerica, MA) in a sterile 35mm petri dish (Falcon) with 3 ml of pro- proliferation medium[10]: low glucose DMEM(Gibco) supplemented with 10% FBS (Biowest) and 100U/mL penicillin/streptomycin (Gibco).

To capture the Dbp rhythm in wounded skin explants, we repeated the same experiment with the explants from K14-cre Dbp:Luc animals. In addition to the explants wounded at the end of T25 entrainment, a second group of samples were wounded 12 hours after entrainment concluded to give a 12hr shift in injury stimulation. These samples were cultured in the same bicarbonate-buffered recording media for explant recording in a sealed petri dish and recorded in the LumiCycle (Actimetrics) at 37°C after wounding (Fig.4A).

**Figure 3.**
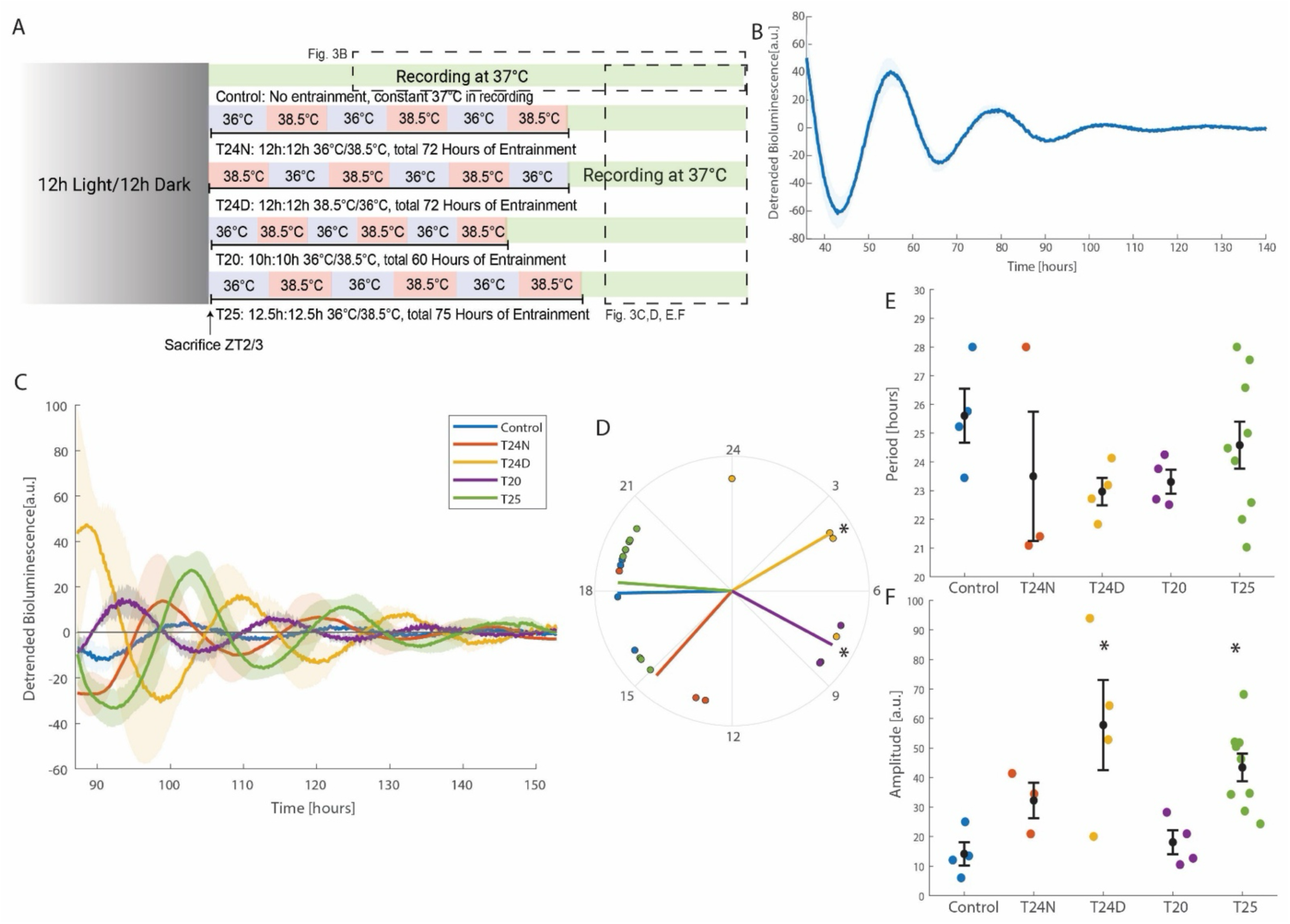
ThermoClock entrained Dbp rhythms in ex vivo murine skin explants. A) Dorsal skin explants of K14-cre Dbp:luc mice were extracted using a biopsy punch after euthanasia. Explants from each animal were cultured on inserts at air-liquid interface and entrained by different temperature cycles on the ThermoClock, with a control sample sent to LumiCycle recording at 37℃ without entrainment. B) Detrended bioluminescence recording of rhythmic control Dbp:luc skin explants (with no entrainment, maintained at 37℃) 36h to 140h after euthanasia (N=4, mean±SEM). C) Detrended bioluminescence recording of rhythmic Dbp:luc skin explants 87h to 150h after euthanasia, starting 12h after last entrainment regime (T25) completed. D, E, F) Phases, periods and amplitudes, respectively, of entrainment groups and controls (fitted from 87h to 167h after euthanasia; N=3, mean±SEM; one-way repeated measures of ANOVA with Dunnett’s post hoc test compared to constant 37, \**p* < 0.05) *Time in Fig.3B,C is time since euthanasia, also the time since the control samples began recording.

**Figure 4.**
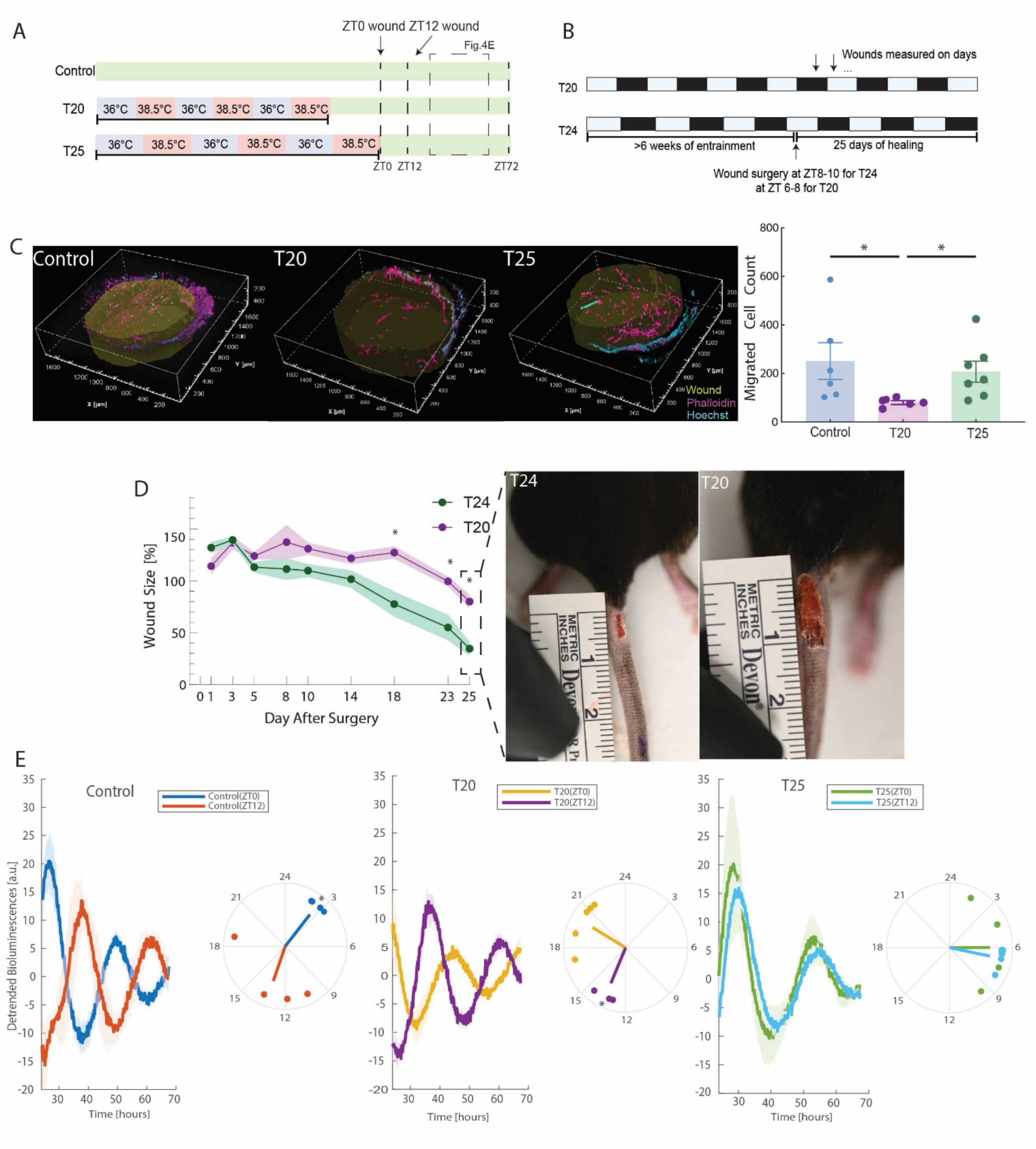
T20 temperature cycle entrainment delays wound healing response ex vivo, similarly to T20 light cycle in vivo disruption, and prior entrainment modulates local oscillator’s response to wounding as a circadian resetting cue. A) Dorsal skin samples were either cultured at 37℃ or entrained by T20, T25 temperature T-cycles before wounded at two different time points (ZT0 or ZT12). Explants were allowed to heal at 37℃ after wounding; samples for proliferation evaluation were collected 72h after wounding. B) Mice were housed under either T24 light cycles (12h:12h light/dark) or T20 light cycles(10h:10h light/dark) for more than 6 weeks before surgery, and wound closure was monitored for 25 days. C) 3D- reconstruction of immunofluorescent fibrin hydrogel inside the explant wounds, and quantified cell migration into the hydrogel (N≥9, mean±SEM; one-way ANOVA with Tukey’s post hoc test, \**p* < 0.05). D) In vivo tail-wound closure under T24 and T20 light-cycle conditions. Wound size is shown as percentage of the original wound area over 25 days after surgery, with representative wound images from day 25. T20-housed mice showed delayed wound closure compared with T24 controls (N≥4, are mean ± SEM; multiple t-tests with Holm-Šídák correction, \**p* < 0.05). E) Detrended bioluminescence recording and fitted phases of rhythmic Dbp:Luc skin explants from hour 24 to hour 68. (N≥3, are mean ± SEM; student’s t-tests on phases, \**p* < 0.05) *Time in Fig.4E is time since end of T25 entrainment/ZT0 wound.

### Bioluminescence Recordings and Analysis

Sealed sample dishes of cell culture or tissue explants were placed in a LumiCycle (Actimetrics) kept at 37°C in an air incubator, without additional humidity or CO_2_[34]. Bioluminescence of individual samples was recorded for 1min every 10 min over at least 7 days unless otherwise noted. At the end of recording, samples were fixed with 4% paraformaldehyde (PFA, Invitrogen) overnight at 4°C.

Bioluminescence recordings were analyzed in a custom MATLAB (MathWorks, Natick, MA) script based on methods in previously published studies[33], [35]. For all cell culture recordings, the first 24 hours of recording was discarded, followed by detrending via removing 24h moving window average. Because of the 24h average method, 12h of data from hour 24 to hour 36 was excluded to avoid edge effects. The first 80 hours of detrended time series (i.e., hour 36 to 116 from start of recording) was then fitted to a damped cosine function with a linear baseline and a constant offset:

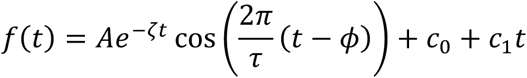

where *A* is the oscillation amplitude, *ζ* the damping rate, *τ* the period, and *φ* the phase. Nonlinear least squared analysis was used to perform fit and the fit quality is assessed based on the coefficient of determination (*R^2^*).

For explant recordings, similar detrending and fitting methods were applied as described above. However, for inter-condition comparisons, we only considered data from 75 hours after euthanasia, starting at the end of T25 entrainment (i.e., the longest entrainment cycle). For wounded explants, only 44h of the detrended time series (from hour 24 to hour 68 after the end of T25 entrainment) was fitted. Additionally, the rhythmicity p-value of each sample was calculated by applying Fast Fourier Transform (FFT) spectral analysis on detrended data to identify the power of frequencies between 1/20 (20h period) to 1/30 (30h period). Background spectral powers were fitted to a log-log linear regression, and a z-score based rhythmicity score, as well as a *p*-value were calculated for testing the hypothesis that the power associated with the circadian period is significantly larger than background noise. The data of individual samples was only included in further parameter analysis if rhythmicity *p*-value ≤ 0.05, and *R^2^* of regression fitting ≥0.85.

A minimum of three independent biological replicates was required for in vitro experiments. For ex vivo studies, analysis required at least three rhythmic samples obtained from

three independent animals (N ≥ 3). Experimental conditions generated from the same cell preparation and multiple skin samples collected from the same animal were treated as repeated measures. Data were analyzed using a one-way repeated-measures ANOVA, followed by Šídák’s or Dunnett’s post hoc test for multiple comparisons, as appropriate.

### Histological and Immunostaining Analysis

Unwounded skin samples were embedded in optimal cutting temperature (OCT) compound (Thermo Fisher Scientific) and cryosectioned at a thickness of 8 µm. A TUNEL assay (Roche) was performed on the tissue sections according to the manufacturer’s instructions. The relationship between the percentage of TUNEL-positive cells and the rhythmicity score was evaluated using simple linear regression using GraphPad Prism (GraphPad Software, Boston MA). TUNEL positivity was compared across experimental conditions using a one-way repeated-measures ANOVA followed by Dunnett’s post hoc test for multiple comparisons against the control condition.

Wounded skin samples from wildtype animals were stained directly with Phalloidin-iFluor 647 Reagent (1:1000, Abcam) and DAPI (1:10000, Sigma-Aldrich). Z-stacked images were captured by NIKON AXR-NSPARC point scanning confocal microscopy across 250 µM thickness. 3D reconstruction and quantification were performed on NIKON NIS-element General Analysis 3 (GA3).

### In Vivo Tail Skin Wounding Assay

Wildtype C57BL/6 animals 2-3 months of age, mixed genders, were group housed under 12h:12h light/dark (T24 control) or under 10h:10h light/dark (T20 circadian disrupted) for 6 weeks prior to surgery using Circadian Cabinets (Phenome Technologies, Lincolnshire, IL). For T24 control animals, surgery occurred at ZT8-10, for T20 circadian disrupted animals at ZT 6-8, both around the midpoint of their respective light-on period. 3.0mmx10.0mm (width by length) full thickness wounds were made on the tail of the animals under analgesic and anesthesia. Following the surgery, wounds were measured manually every 2-4 days up to day 25.

## Results

### ThermoClock temperature cycles enhance reporter cell line entrainment compared with incubator-based cycles

We designed the ThermoClock as a low-cost and easy-to-manufacture temperature control device that would be particularly suitable for temperature entrainment and synchronization in circadian studies (Fig.1A). While the system only provides temperature control, it is fully compatible to use inside a standard CO_2_ incubator under 95% humidity to achieve optimal cell growth conditions. The system has also been used successfully under hypoxic conditions in a CO_2_ incubator with low O_2_ settings balanced with N_2_.

The ThermoClock achieved setpoint temperatures more rapidly than a programmable incubator allowing for a user-input temperature schedule (Fig.1B) [29]. Our previous publication demonstrated that ThermoClock heats up to setpoint temperatures within 5 mins during heating and passively cools to setpoints within 20-40 mins, while it takes the programmable incubator over an hour to accomplish each process [29]. We validated the technical temperature control aspects robustly in our previous publication – including the demonstration that all 5 plates can be programmed with individual temperature profiles – hence here we focus on validating the application of ThermoClock in biological applications.

We first used the ThermoClock to temperature entrain circadian reporter cell lines in 2D culture: Bmal1:Luc and Per2:Luc U2OS[33]. As commonly used model cell lines, U2OS rhythms are often synchronized with a 2-hour dexamethasone (Dex) pulse [33], [36].

Synchronized U2OS cultures show higher bioluminescence amplitudes, which implies that the circadian rhythms of cells are more in phase with each other. To compare the ThermoClock against a commercially available product, we also included the same incubator that allows user-input temperature schedules mentioned above: both instruments were programmed to entrain cells with 3 cycles of 12h:12h 36°C/38.5°C, a temperature profile that has been previously demonstrated to entrain peripheral cultures effectively [23]. A dexamethasone-synchronized group and a constant 37°C culture group serve as the reference conditions (Fig.2A). All the reference conditions’ samples were plated at the same time as the entrainment groups but instead maintained at 37°C for three days; the start of dexamethasone pulse was aligned with the end of the temperature entrainments (Fig.2A). Following these steps, bioluminescence data was then captured for seven days at 37°C, with all samples sealed and maintained in sodium bicarbonate-buffered recording media since the recording device is maintained at atmospheric conditions without added humidity.

Non-linear regression outputs revealed significantly increased Bmal1 and Per2 amplitude in samples entrained by the ThermoClock (Fig 2B, C, D, E) compared to no treatment negative controls maintained at 37°C. To our surprise, the incubator-based entrainment, while employing the same temperature trajectory, did not increase either Bmal1 or Per2 amplitudes (Fig. 2D, E) significantly. Bmal1 and Per2 rhythms remained anti-phase after ThermoClock entrainment and no significant changes to the averaged periods or damping rates were observed (Fig. 2D, F; Supplementary Fig.1). Periods of 3 consecutive cycles also didn’t show significant deviation to control conditions (Supplementary Fig.1). However, an approximately four-hour phase delay in Bmal1 and Per2 rhythms were observed in both temperature entrainment groups compared to controls (Fig. 2E, G). The comparison between ThermoClock and the incubators suggests a new insight that strength of temperature cycles as a zeitgeber is not only dependent on magnitude of the temperature change but also impacted by the waveform of the thermal oscillation, specifically on the ramp time to setpoint temperatures.

### K14-cre Dbp:Luc reporter skin explants could be entrained to circadian temperature cycles and T-cycles

3D cultures serve as important biological models capturing structural, cell-extracellular matrix interactions, and the capacity for multi-cellular complexity [37], [38]. The most complex of these, tissue explants, can be used directly to test treatments [39], [40]; to probe mechanisms [41], [42]; or to provide important insights into creating more physiologically relevant bioengineered systems [43], [44]. Patient-derived tissue explants are also often used as personalized disease models[39]. However, like cell cultures, circadian rhythms in peripheral tissue explants damp rapidly as they detach from the endogenous time-keeping system [17], [18], [19]. We thereby chose to challenge ThermoClock to produce rhythms in this complex 3D tissue environment.

To explore maintaining circadian rhythms in tissue explants via temperature entrainment using the ThermoClock, we entrained skin tissues extracted from albino C57BL/6 animals expressing the Dbp:Luc reporter in keratinocytes (Fig.3A). Dbp is known to be rhythmically expressed in different organs and tissues [45], [46], [47]. In this line of animals, the luciferase reporter of the mouse Dbp is Cre-recombinase-dependent to turn on the bioluminescence reporter specifically in keratinocytes [31], [32]. To avoid unintentional rhythm resets on the cultures, the explants were entrained directly in sealed dishes with bicarbonate-buffered recording media and transferred to the LumiCycle as soon as their respective entrainments were completed. Four entrainment schedules were employed simultaneously on the ThermoClock: T24 Diurnal (T24D, 12h:12h 38.5°C/36°C), T24 Nocturnal (T24N, 12h:12h 36°C/38.5°C), T20 (10h:10h 36°C/38.5°C), T25 (12.5h:12.5h 36°C/38.5°C; Fig.3A). The opposing T24 conditions were included to confirm that entrainment of peripheral tissues such as skin results in oscillations in phase with the entrainment signal. The non-24-hour T-cycles T20 and T25 were included as potential ways to induce circadian disruption in vitro, a technique which has not been previously explored.

Bioluminescence data were included in analysis only if the detrended data passed the FFT-based rhythmicity test and fit quality threshold (*p* > 0.05, R^2^>0.85; Supplementary Fig.2). TUNEL assay showed no significant correlation between the level of cell death and entrainment approaches, confirming that ThermoClock-based temperature entrainment does not increase cell death (Supplementary Fig. 3). Under no entrainment and cultured at 37℃, Dbp:Luc skin explants maintained robust but rapidly damping oscillation for approximately the first 90h in ex vivo culture (Fig.3B). Recording of entrained explants began at 75h post-euthanasia and tissue collection. Unlike previous experiments, no transient period was removed from these samples, as we expect the luciferase-luciferin ratio has equilibrated before recording. After ThermoClock entrainment, Dbp rhythms of the skin explants were robust, and explant rhythms were able to adopt the phase of temperature cycles, as shown by the ∼12h difference in phase between the T24D and the T24N groups (Fig.3C, D). These observations are consistent with previous studies on peripheral tissue (i.e., lung and liver) entrainment[23], [28], [48].

Next, we investigated temperature T-cycle entrainment in explants. While neither T20 nor T25 entrainment groups resulted in changes in Dbp periods (Fig. 3E), T20-entrained samples exhibited damped amplitudes similar to the non-entrained control, and a ∼10h phase advance compared to control, T24 nocturnal, and T25 samples. T25 entrainment, by contrast, produced the opposite outcome — higher amplitude oscillations with no phase shift compared to control, and the highest proportion of rhythmic samples across all groups (Fig. 3D, F; Supplementary Figure 2). These findings suggest that T25 acts as a strong synchronizer to Dbp rhythm in skin. Meanwhile, T20 induced partial or conflicted entrainment, with some cells adjusting to the entrainment phase while the majority cannot follow it. It is worth noting that all the above entrainments were performed simultaneously by a ThermoClock contained within one incubator. Beyond automating the process, our device significantly reduced equipment, space, and energy expenditure compared to traditional approaches, while maximizing experimental throughput, and limiting the potential variations introduced by varied devices or non-parallelized studies.

### T20 entrained skin explants showed reduced cell migration in ex vivo wounding assay consistent with in vivo observation

While ex vivo peripheral systems show more flexible entrainment to extreme T-cycles, few have investigated whether there is an entrainment after-effect in cellular function. T20 light entrainment (10h:10h light/dark) is a classic paradigm for modeling circadian disruption; it induces physiological changes including increased weight gain, altered immune response, and altered cognitive functions[24], [25]. Although animals living in T20 conditions do not entrain to the 20-hour period in their locomotor activity or core body temperature, the physiological changes suggest that the misalignment between the endogenous clock and the external zeitgeber cycle induces stress to the overall system. This begs the question: does applying T20 temperature entrainment on isolated peripheral clock tissues mimic the pressure imposed by T20 LD schedules in vivo? ThermoClock offers a robust way of testing this that has not been previously explored as an approach.

Here, we developed an ex vivo skin wound healing model – skin explants were wounded at the end of entrainment by a biopsy punch and the created “wound” was filled with a fibrin hydrogel to mimic a blood clot and provide a matrix for cell migration. Significantly less cell migration was observed in samples entrained by T20 temperature cycles compared to T25 entrained samples and controls (Fig.3B). This observation supports the idea that skin cells retain acute memories from ex vivo temperature entrainment and their cellular functions are impacted by temperature T-cycle entrainments.

To see how this compares to in vivo behavior, we performed a tail wound healing assay comparing animals under normal T24 circadian light conditions (12h:12h light/dark) and T20 circadian disruption (10h:10h light/dark; Fig.4B). We chose the tail wound model to reduce the impact from prolonged surgery and anesthesia time on the animals’ rhythms, as well as to avoid the confounding factor of skin contracting around the wound[49], [50]. A significant difference in wound size was observed towards the end of the observed window on days 18, 23 and 25. On day 25, there remained a near two-fold gap between T24 (34.6% of original wound remained) and T20 (80% of original wound remained) animals (Fig.4C), which, remarkably, is very close to the difference observed in our ThermoClock entrained T20 versus T25 samples ex vivo.

Collectively, these results suggested that our ThermoClock entrained explants were able to recapitulate at least part of the perturbed healing response to injury under circadian disruption.

### Entrainments impacted sensitivity of explants to rhythm reset by wounding

Previous studies have shown that dissection can reset a tissue’s circadian phase from the endogenous in vivo rhythm, and the resetting sensitivity is increased in animals entrained by T20 light cycles [51]. This inspired us to investigate if injury, like dissection, can reset peripheral clocks; and if explants entrained by temperature cycles will show different sensitivities to resetting cues. To answer these questions, we monitored the Dbp rhythm on the K14-cre Dbp:Luc explants that followed the same ex vivo wounding protocol (Fig.3C, Supplementary Fig. 4). All samples – regardless of pre-injury conditions – were separated into two groups wounded 12h apart. Non-entrained samples exhibited near anti-phase oscillation following the injuries, suggesting that wounding can indeed act as a synchronizer. However, this phase-resetting response is dependent on the pre-injury entrainment condition as we observed a ∼6h phase difference in the T20 entrained wounds, and no significant phase adjustment was observed in the T25 samples (Fig.4D). These results indicate that wounding can reset Dbp rhythms and potentially the local oscillator in skin explants; and, the sensitivity of the tissue to injury-induced resetting is affected by state of the clock pre-injury as set by ex vivo temperature entrainments.

In this work, a more robust entrainment (T25) lead to a resistance to resetting of the clock in response to injury, while a disrupted clock (T20) resulted in a phase shift.

## Discussion

In this study, we present evidence of the biological efficacy of ThermoClock, a low-cost platform that enables controlled, reproducible temperature entrainment of both 2D cell lines and 3D tissue explants. Temperature is a physiologically potent zeitgeber for peripheral clocks, yet it remains underexplored due to a lack of accessible tools – current approaches of using programmable incubators are costly and have a large physical footprint, limiting experimental throughput and multiplexing capacity. Made with off-the-shelf electronics and an affordable microcontroller, the ThermoClock can be easily integrated into existing standard lab setups to treat cultures with different temperature cycles. Another key advantage of the ThermoClock is that it enables precise control over temperature waveform parameters, including amplitude, period, phase, and ramp time to setpoint temperatures (which is tuned by varying the control parameters of ThermoClock). As ThermoClock’s heat source is directly adjacent to the cells, the temperature profile can be transferred much more quickly than trying to heat/cool an entire incubator. This level of control allowed us to directly compare entrainment outcomes across thermal profiles, which would be inefficient and difficult to do using conventional incubator-based methods. As revealed by the Bmal1 and Per2 rhythms in U2OS following entrainment by different devices, the shorter ramp times of ThermoClock versus an incubator resulted in a higher amplitude of the entrained rhythms (Fig. 2). These results suggest that the waveform of the temperature input – not only their absolute amplitude and period – is an important parameter in clock entrainment.

Currently, considerable effort in biomedical engineering focuses on developing physiologically relevant culture systems, including organoids, patient-specific explants and tissue-on-chip platforms; yet circadian regulation is almost universally neglected in these systems [20], [21]. Here, we challenged ThermoClock on skin explants extracted from animals with keratinocyte-specific Dbp:luc reporter and entrained samples simultaneously through different temperature programs. This capacity to multiplex up to 5 individual entrainment schedules on one ThermoClock device significantly increased the flexibility in study design and lowered experimental variations. As ThermoClock sits on a single incubator shelf, multiple devices can be used if more temperature profiles are needed. We demonstrated that ThermoClock entrainment can 1) extend the circadian rhythms in these explants so that time-of-day differences can be recapitulated under ∼24h temperature cycles, 2) induce circadian disruption in explants tissues to assess specific functional changes using non-circadian T-cycles, like the T20. In this study, our parallel T20 ex vivo temperature-entrained skin wounds and in vivo light-entrained wounds showed similar attenuation in wound healing response. While chemical approaches to culture synchronization in vitro predominantly function to force the clock into alignment or target a single component of the clock, temperature T-cycles impose a conflict between the endogenous oscillator and an external zeitgeber, making temperature entrainment more analogous to in vivo light cycle disruption. Understanding that wound healing is a multi-cellular and multi-stage process, these findings still suggest that mimicking disruption with temperature in an isolated ex vivo system can capture the delayed healing. Considering the significantly shortened timeframe (3 days of entrainment ex vivo versus 6 weeks in vivo), we think this can serve as a powerful first-pass screening for therapeutics targeting physiological dysfunction under circadian disruption. Furthermore, these in vitro model systems remove crosstalk complexity by allowing insights into specific questions. Ongoing studies in our lab are looking to engineer multicellular systems cultured under different temperature cycles to more fully recapitulate the healing dynamics in different scenarios.

By coupling circadian rhythm recording with a functional wound-healing assay, we further showed that the state of the clock can shape its response to perturbations such as injury. Non-entrained, T20-disrupted, and T25-entrained explants displayed distinct sensitivities to wound-induced phase resetting, suggesting that prior entrainment history alters the sensitivity of the tissue clock. This observation is consistent with previous modeling studies showing that coupling among cellular clocks can influence the rigidity of local oscillators and the magnitude of their response to resetting cues [35], [52], [53]. Although non-entrained explants did not show reduced migration, we observed a reset of the Dbp rhythm by the injury. We attributed this to a synchronizing effect of the injury; notably, T25 entrained samples with robust clocks resisted this synchronization signal. This is an important insight for in vitro biomedical models: maintaining cultures in the absence of a robust circadian clock may fail to capture not only the time-of-day variations, but also normal repeated experimental perturbations (e.g., injury, mechanical or pharmacological stimulations, media change) could be triggering unintended clock resets and alter downstream tissue responses.

In conclusion, the ThermoClock makes precise thermal regulation accessible and practical for in vitro model systems. Programmable incubators are expensive, occupy large footprints, and limit throughput to one temperature profile per unit. The ThermoClock delivers up to five independent programs simultaneously from a single incubator shelf at a fraction of the cost, making it a versatile tool not only for circadian biologists but for any researcher requiring controlled thermal modulation—including studies of heat shock responses and thermal regulation in immunity. This work also establishes temperature T-cycles as a promising strategy for modeling circadian disruption in vitro—a capability largely absent to date. The functional alterations we observe suggest that this represents physiologically meaningful disruption with measurable downstream effects. As complex in vitro platforms increasingly enter the therapeutic development pipeline, and we strive to faithfully recapitulate physiology with these models, the ability to model both healthy and disrupted circadian rhythms will be essential for capturing this temporal dimension of human disease.

## Acknowledgement

Research reported in this publication was supported by the National Institute of General Medical Sciences of the National Institutes of Health under award number R35GM147272 and R15GM126545, and a UMass Amherst Interdisciplinary Grant. We also want to acknowledge the imaging and tissue processing support from UMass Amherst Light Microscopy and Histology Core Facilities.

We would like to honor the memory and contribution of our co-author, Bhavna Kalyanaraman, Ph.D., who passed away prior to the publication of this work.

## References

[1] D. Montaigne et al., “Daytime variation of perioperative myocardial injury in cardiac surgery and its prevention by Rev-Erbα antagonism: a single-centre propensity-matched cohort study and a randomised study,” The Lancet, vol. 391, no. 10115, pp. 59–69, Jan. 2018, doi: 10.1016/S0140-6736(17)32132-3.

[2] A. B. Fishbein, K. L. Knutson, and P. C. Zee, “Circadian disruption and human health,” J. Clin. Invest., vol. 131, no. 19, p. e148286, Oct. 2021, doi: 10.1172/JCI148286.

[3] M. F. Gonzalez-Aponte et al., “Circadian variation in MGMT promoter methylation and expression predicts sensitivity to temozolomide in glioblastoma,” J. Neurooncol., vol. 176, no. 1, p. 36, Jan. 2026, doi: 10.1007/s11060-025-05242-3.

[4] G. L. Pearson et al., “Time of day alters olfactory bulb immune state with ramifications for intranasal inflammatory challenge,” Cell Rep., vol. 45, no. 4, p. 117133, Apr. 2026, doi: 10.1016/j.celrep.2026.117133.

[5] M. I. Printezi et al., “Toxicity and efficacy of chronomodulated chemotherapy: a systematic review,” Lancet Oncol., vol. 23, no. 3, pp. e129–e143, Mar. 2022, doi: 10.1016/S1470-2045(21)00639-2.

[6] A. Kalsbeek et al., “SCN Outputs and the Hypothalamic Balance of Life,” J. Biol. Rhythms, vol. 21, no. 6, pp. 458–469, Dec. 2006, doi: 10.1177/0748730406293854.

[7] Y. Xie et al., “New Insights Into the Circadian Rhythm and Its Related Diseases,” Front. Physiol., vol. 10, p. 682, 2019, doi: 10.3389/fphys.2019.00682.

[8] H. Jacob, A. M. Curtis, and C. J. Kearney, “Therapeutics on the clock: Circadian medicine in the treatment of chronic inflammatory diseases,” Biochem. Pharmacol., vol. 182, p. 114254, Dec. 2020, doi: 10.1016/j.bcp.2020.114254.

[9] S. L. Chellappa, N. Vujovic, J. S. Williams, and F. A. J. L. Scheer, “Impact of Circadian Disruption on Cardiovascular Function and Disease,” Trends Endocrinol. Metab., vol. 30, no. 10, pp. 767–779, Oct. 2019, doi: 10.1016/j.tem.2019.07.008.

[10] N. P. Hoyle et al., “Circadian actin dynamics drive rhythmic fibroblast mobilization during wound healing,” Sci. Transl. Med., vol. 9, no. 415, p. eaal2774, Nov. 2017, doi: 10.1126/scitranslmed.aal2774.

[11] E. J. Cable, K. G. Onishi, and B. J. Prendergast, “Circadian rhythms accelerate wound healing in female Siberian hamsters,” Physiol. Behav., vol. 171, pp. 165–174, Mar. 2017, doi: 10.1016/j.physbeh.2016.12.019.

[12] S. M. Schmid, M. Hallschmid, and B. Schultes, “The metabolic burden of sleep loss,” Lancet Diabetes Endocrinol., vol. 3, no. 1, pp. 52–62, Jan. 2015, doi: 10.1016/S2213-8587(14)70012-9.

[13] X. Xiong, T. Hu, Z. Yin, Y. Zhang, F. Chen, and P. Lei, “Research advances in the study of sleep disorders, circadian rhythm disturbances and Alzheimer’s disease,” Front. Aging Neurosci., vol. 14, p. 944283, Aug. 2022, doi: 10.3389/fnagi.2022.944283.

[14] M. D. Ruben, D. F. Smith, G. A. FitzGerald, and J. B. Hogenesch, “Dosing time matters,” Science, vol. 365, no. 6453, pp. 547–549, Aug. 2019, doi: 10.1126/science.aax7621.

[15] C. R. Cederroth et al., “Medicine in the Fourth Dimension,” Cell Metab., vol. 30, no. 2, pp. 238–250, Aug. 2019, doi: 10.1016/j.cmet.2019.06.019.

[16] E. Esposito et al., “Potential circadian effects on translational failure for neuroprotection,” Nature, vol. 582, no. 7812, pp. 395–398, Jun. 2020, doi: 10.1038/s41586-020-2348-z.

[17] Y. Li, Y. Shan, R. V. Desai, K. H. Cox, L. S. Weinberger, and J. S. Takahashi, “Noise-driven cellular heterogeneity in circadian periodicity,” Proc. Natl. Acad. Sci., vol. 117, no. 19, pp. 10350–10356, May 2020, doi: 10.1073/pnas.1922388117.

[18] D. K. Welsh, S.-H. Yoo, A. C. Liu, J. S. Takahashi, and S. A. Kay, “Bioluminescence Imaging of Individual Fibroblasts Reveals Persistent, Independently Phased Circadian Rhythms of Clock Gene Expression,” Curr. Biol., vol. 14, no. 24, pp. 2289–2295, Dec. 2004, doi: 10.1016/j.cub.2004.11.057.

[19] Y. Li et al., “Epigenetic inheritance of circadian period in clonal cells,” eLife, vol. 9, p. e54186, May 2020, doi: 10.7554/eLife.54186.

[20] J.-M. Fustin, M. Li, B. Gao, Q. Chen, T. Cheng, and A. G. Stewart, “Rhythm on a chip: circadian entrainment *in vitro* is the next frontier in body-on-a chip technology,” Curr. Opin. Pharmacol., vol. 48, pp. 127–136, Oct. 2019, doi: 10.1016/j.coph.2019.09.005.

[21] K. J. Cyr, O. M. Avaldi, and J. P. Wikswo, “Circadian hormone control in a human-on-a-chip: In vitro biology’s ignored component?,” Exp. Biol. Med., vol. 242, no. 17, pp. 1714– 1731, Nov. 2017, doi: 10.1177/1535370217732766.

[22] C. Dibner, U. Schibler, and U. Albrecht, “The Mammalian Circadian Timing System: Organization and Coordination of Central and Peripheral Clocks,” Annu. Rev. Physiol., vol. 72, no. 1, pp. 517–549, Mar. 2010, doi: 10.1146/annurev-physiol-021909-135821.

[23] E. D. Buhr, S.-H. Yoo, and J. S. Takahashi, “Temperature as a universal resetting cue for mammalian circadian oscillators,” Science, vol. 330, no. 6002, pp. 379–385, Oct. 2010, doi: 10.1126/science.1195262.

[24] I. N. Karatsoreos, S. Bhagat, E. B. Bloss, J. H. Morrison, and B. S. McEwen, “Disruption of circadian clocks has ramifications for metabolism, brain, and behavior,” Proc. Natl. Acad. Sci. U. S. A., vol. 108, no. 4, pp. 1657–1662, Jan. 2011, doi: 10.1073/pnas.1018375108.

[25] F. Fuchs et al., “Delaying circadian sleep phase under ultradian light cycle causes time-of-day-dependent alteration of cognition and mood,” Sci. Rep., vol. 13, no. 1, p. 20313, Nov. 2023, doi: 10.1038/s41598-023-44931-9.

[26] A. Balsalobre et al., “Resetting of Circadian Time in Peripheral Tissues by Glucocorticoid Signaling,” Science, vol. 289, no. 5488, pp. 2344–2347, Sep. 2000, doi: 10.1126/science.289.5488.2344.

[27] A. Balsalobre, F. Damiola, and U. Schibler, “A Serum Shock Induces Circadian Gene Expression in Mammalian Tissue Culture Cells,” Cell, vol. 93, no. 6, pp. 929–937, Jun. 1998, doi: 10.1016/S0092-8674(00)81199-X.

[28] S. A. Brown, G. Zumbrunn, F. Fleury-Olela, N. Preitner, and U. Schibler, “Rhythms of Mammalian Body Temperature Can Sustain Peripheral Circadian Clocks,” Curr. Biol., vol. 12, no. 18, pp. 1574–1583, Sep. 2002, doi: 10.1016/S0960-9822(02)01145-4.

[29] K. K. Zhang, D. Locurto, M. Belden, M. Price, C. J. Kearney, and M. E. Huber, “Around the ThermoClock: A precision automated temperature control system for Ex Vivo circadian studies,” MethodsX, vol. 15, p. 103717, Dec. 2025, doi: 10.1016/j.mex.2025.103717.

[30] E. D. Buhr and J. S. Takahashi, “Molecular components of the Mammalian circadian clock,” Handb. Exp. Pharmacol., no. 217, pp. 3–27, 2013, doi: 10.1007/978-3-642-25950-0_1.

[31] C. B. Smith et al., “Cell-Type-Specific Circadian Bioluminescence Rhythms in Dbp Reporter Mice,” J. Biol. Rhythms, vol. 37, no. 1, pp. 53–77, Feb. 2022, doi: 10.1177/07487304211069452.

[32] V. Vasioukhin, L. Degenstein, B. Wise, and E. Fuchs, “The magical touch: Genome targeting in epidermal stem cells induced by tamoxifen application to mouse skin,” Proc. Natl. Acad. Sci., vol. 96, no. 15, pp. 8551–8556, Jul. 1999, doi: 10.1073/pnas.96.15.8551.

[33] H.-H. Lin, K. L. Robertson, H. A. Bisbee, and M. E. Farkas, “Oncogenic and Circadian Effects of Small Molecules Directly and Indirectly Targeting the Core Circadian Clock,” Integr. Cancer Ther., vol. 19, Jan. 2020, doi: 10.1177/1534735420924094.

[34] S. Yamazaki and J. S. Takahashi, “Real-Time Luminescence Reporting of Circadian Gene Expression in Mammals,” in Methods in Enzymology, vol. 393, Elsevier, 2005, pp. 288–301. doi: 10.1016/S0076-6879(05)93012-7.

[35] T. L. Leise, C. W. Wang, P. J. Gitis, and D. K. Welsh, “Persistent Cell-Autonomous Circadian Oscillations in Fibroblasts Revealed by Six-Week Single-Cell Imaging of PER2::LUC Bioluminescence,” PLoS ONE, vol. 7, no. 3, p. e33334, Mar. 2012, doi: 10.1371/journal.pone.0033334.

[36] C. Vollmers, S. Panda, and L. DiTacchio, “A high-throughput assay for siRNA-based circadian screens in human U2OS cells,” PloS One, vol. 3, no. 10, p. e3457, 2008, doi: 10.1371/journal.pone.0003457.

[37] K. Duval et al., “Modeling Physiological Events in 2D vs. 3D Cell Culture,” Physiology, vol. 32, no. 4, pp. 266–277, Jul. 2017, doi: 10.1152/physiol.00036.2016.

[38] R. Xie et al., “A comprehensive review on 3D tissue models: Biofabrication technologies and preclinical applications,” Biomaterials, vol. 304, p. 122408, Jan. 2024, doi: 10.1016/j.biomaterials.2023.122408.

[39] I. R. Powley et al., “Patient-derived explants (PDEs) as a powerful preclinical platform for anti-cancer drug and biomarker discovery,” Br. J. Cancer, vol. 122, no. 6, pp. 735–744, Mar. 2020, doi: 10.1038/s41416-019-0672-6.

[40] F. Rancan et al., “Evaluation of Drug Delivery and Efficacy of Ciprofloxacin-Loaded Povidone Foils and Nanofiber Mats in a Wound-Infection Model Based on Ex Vivo Human Skin,” Pharmaceutics, vol. 11, no. 10, p. 527, Oct. 2019, doi: 10.3390/pharmaceutics11100527.

[41] L. M. Prolo, J. S. Takahashi, and E. D. Herzog, “Circadian Rhythm Generation and Entrainment in Astrocytes,” J. Neurosci., vol. 25, no. 2, pp. 404–408, Jan. 2005, doi: 10.1523/JNEUROSCI.4133-04.2005.

[42] J.-C. Grivel and L. Margolis, “Use of human tissue explants to study human infectious agents,” Nat. Protoc., vol. 4, no. 2, pp. 256–269, 2009, doi: 10.1038/nprot.2008.245.

[43] R. Way, H. Templeton, D. Ball, M.-H. Cheng, S. A. Tobet, and T. Chen, “A microphysiological system for studying barrier health of live tissues in real time,” Commun. Eng., vol. 3, no. 1, p. 142, Oct. 2024, doi: 10.1038/s44172-024-00285-2.

[44] B. Lakeh and A. Shafiee, “Advancing dermatology with skin equivalents and organoids in pathophysiology and drug testing,” Acta Biomater., vol. 207, pp. 120–130, Nov. 2025, doi: 10.1016/j.actbio.2025.10.008.

[45] P. Fonjallaz, V. Ossipow, G. Wanner, and U. Schibler, “The two PAR leucine zipper proteins, TEF and DBP, display similar circadian and tissue-specific expression, but have different target promoter preferences,” EMBO J., vol. 15, no. 2, pp. 351–362, Jan. 1996.

[46] S. Punia, K. K. Rumery, E. A. Yu, C. M. Lambert, A. L. Notkins, and D. R. Weaver, “Disruption of gene expression rhythms in mice lacking secretory vesicle proteins IA-2 and IA-2β,” Am. J. Physiol.-Endocrinol. Metab., vol. 303, no. 6, pp. E762–E776, Sep. 2012, doi: 10.1152/ajpendo.00513.2011.

[47] R. Zhang, N. F. Lahens, H. I. Ballance, M. E. Hughes, and J. B. Hogenesch, “A circadian gene expression atlas in mammals: Implications for biology and medicine,” Proc. Natl. Acad. Sci., vol. 111, no. 45, pp. 16219–16224, Nov. 2014, doi: 10.1073/pnas.1408886111.

[48] U. Abraham, A. E. Granada, P. O. Westermark, M. Heine, A. Kramer, and H. Herzel, “Coupling governs entrainment range of circadian clocks,” Mol. Syst. Biol., vol. 6, p. 438, Nov. 2010, doi: 10.1038/msb.2010.92.

[49] V. Falanga et al., “Full-thickness wounding of the mouse tail as a model for delayed wound healing: accelerated wound closure in Smad3 knock-out mice,” Wound Repair Regen., vol. 12, no. 3, pp. 320–326, Jun. 2004, doi: 10.1111/j.1067-1927.2004.012316.x.

[50] M. Yampolsky, I. Bachelet, and Y. Fuchs, “Reproducible strategy for excisional skin-wound-healing studies in mice,” Nat. Protoc., vol. 19, no. 1, pp. 184–206, Jan. 2024, doi: 10.1038/s41596-023-00899-4.

[51] T. L. Leise et al., “Recurring circadian disruption alters circadian clock sensitivity to resetting,” Eur. J. Neurosci., vol. 51, no. 12, pp. 2343–2354, Jun. 2020, doi: 10.1111/ejn.14179.

[52] D. K. Welsh, J. S. Takahashi, and S. A. Kay, “Suprachiasmatic Nucleus: Cell Autonomy and Network Properties,” Annu. Rev. Physiol., vol. 72, no. 1, pp. 551–577, Mar. 2010, doi: 10.1146/annurev-physiol-021909-135919.

[53] Y. Yamaguchi et al., “Mice Genetically Deficient in Vasopressin V1a and V1b Receptors Are Resistant to Jet Lag,” Science, vol. 342, no. 6154, pp. 85–90, Oct. 2013, doi: 10.1126/science.1238599.

