## Supplementary Information for "ThermoClock:A Novel Automated Temperature Regulation Device that Can Model Circadian Entrainment and Disruption in 2D and 3D in Vitro Models"

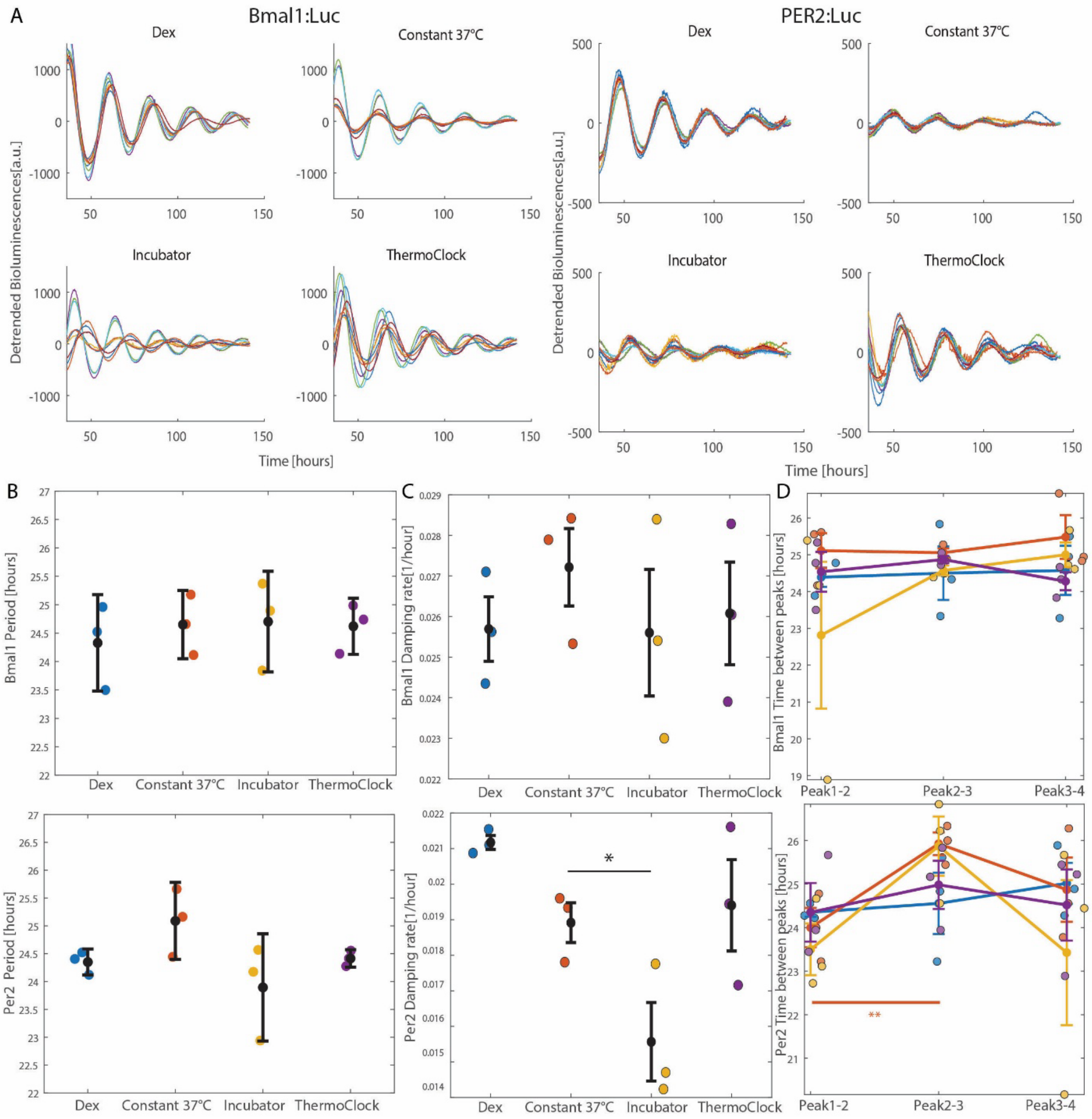


**Supplementary Figure 1. Individual detrended bioluminescence recordings and additional circadian parameters confirm that ThermoClock entrainment does not alter period, damping rate, or rhythm stability.** A) Detrended bioluminescence recordings((under 37°C) of individual Bmal1:Luc and Per2:Luc U2OS samples after dexamethasone synchronization, constant 37°C, incubator or ThermoClock entrainments (N=3, n=3). B) Periods and C) damping rates of Bmal1:Luc and Per2:Luc U2OS cells across synchronization conditions (fitted from 36h to 116h; N=3, mean±SEM; one-way repeated measures ANOVA with Dunnett's post hoc test compared to constant 37°C, *$p < 0.05$). D) Time between consecutive peaks (peak 1–2, peak 2–3, peak 3–4) for Bmal1:Luc and Per2:Luc U2OS cells (fitted from 36h to 116h; N=3, mean±SEM). No significant differences were observed between conditions at each timepoint. Inter-peak intervals did not change significantly, except for Per2:Luc U2OS under the constant 37°C condition (between peak 1–2 and peak 2–3).

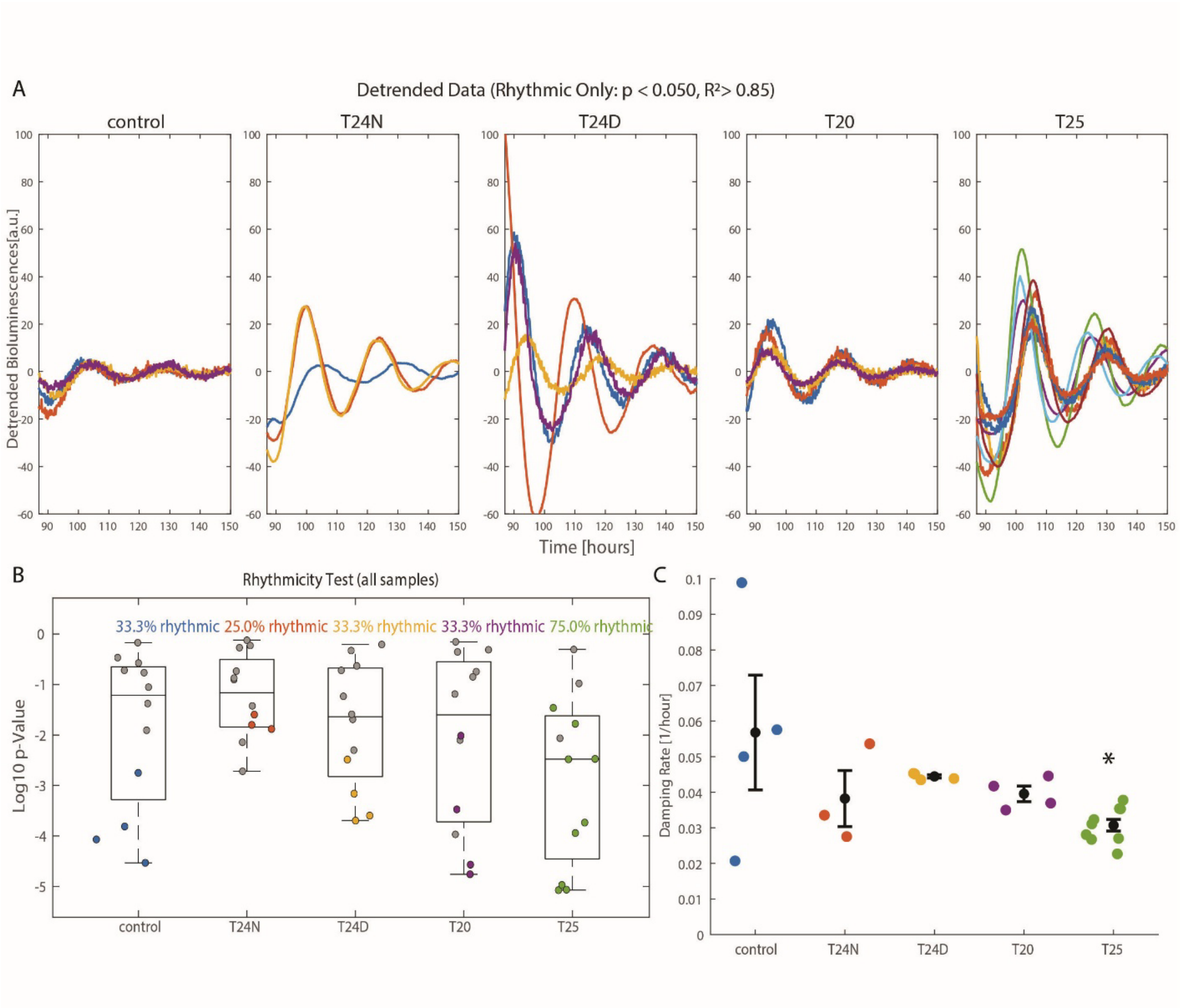


**Supplementary Figure 2. Individual bioluminescence traces, rhythmicity classification, and damping rates of Dbp:Luc skin explants support entrainment findings in Figure 3.** A) Detrended bioluminescence recordings(under 37°C) of individual rhythmic Dbp:Luc skin explants for each condition (control, T24N, T24D, T20, T25) from 87h to 150h after euthanasia (rhythmic samples only: $p < 0.050$, $R^2 > 0.85$; N≥4). B) Rhythmicity test (Log10 p-value) for all samples across entrainment conditions, with percentage of rhythmic samples indicated per group. Samples that failed the rhythmicity and fitness thresholds are greyed. C) Damping rates of rhythmic Dbp:Luc skin explants across entrainment groups and controls (fitted from 87h to 167h after euthanasia; N=3, mean±SEM; one-way repeated measure ANOVA with Dunnett's post hoc test compared to constant 37°C, *$p < 0.050$).

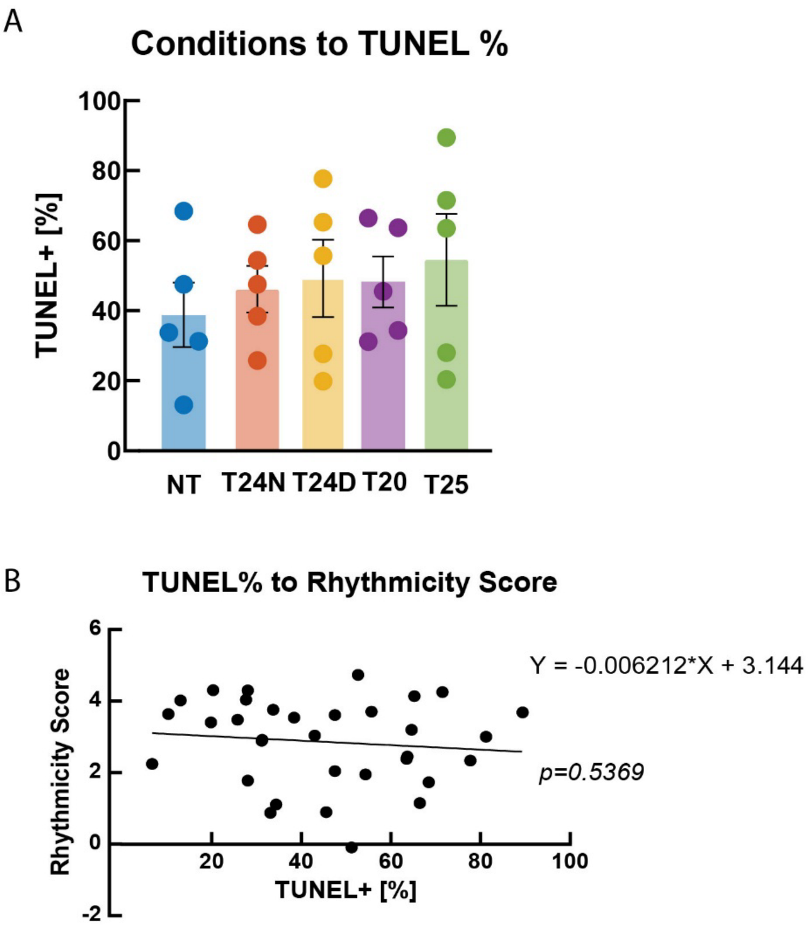


**Supplementary Figure 3. Temperature cycle entrainment does not significantly affect apoptosis in skin explants, and cell death is not correlated with circadian rhythmicity.** A) Percentage of TUNEL-positive cells in Dbp:Luc skin explants across entrainment conditions (NT, T20, T24N, T24D, T25; mean±SEM) at the end of 7 days of recording, 10 days after euthanasia. No significant difference in cell apoptosis was observed between groups (N=5, one-way ANOVA, Tukey's post-hoc test). B) Linear regression of TUNEL-positive percentage against rhythmicity score across all samples. No significant correlation was found between cell death and rhythmicity score ($p$ = 0.5369), indicating that differences in circadian parameters between conditions are not driven by differential tissue viability.

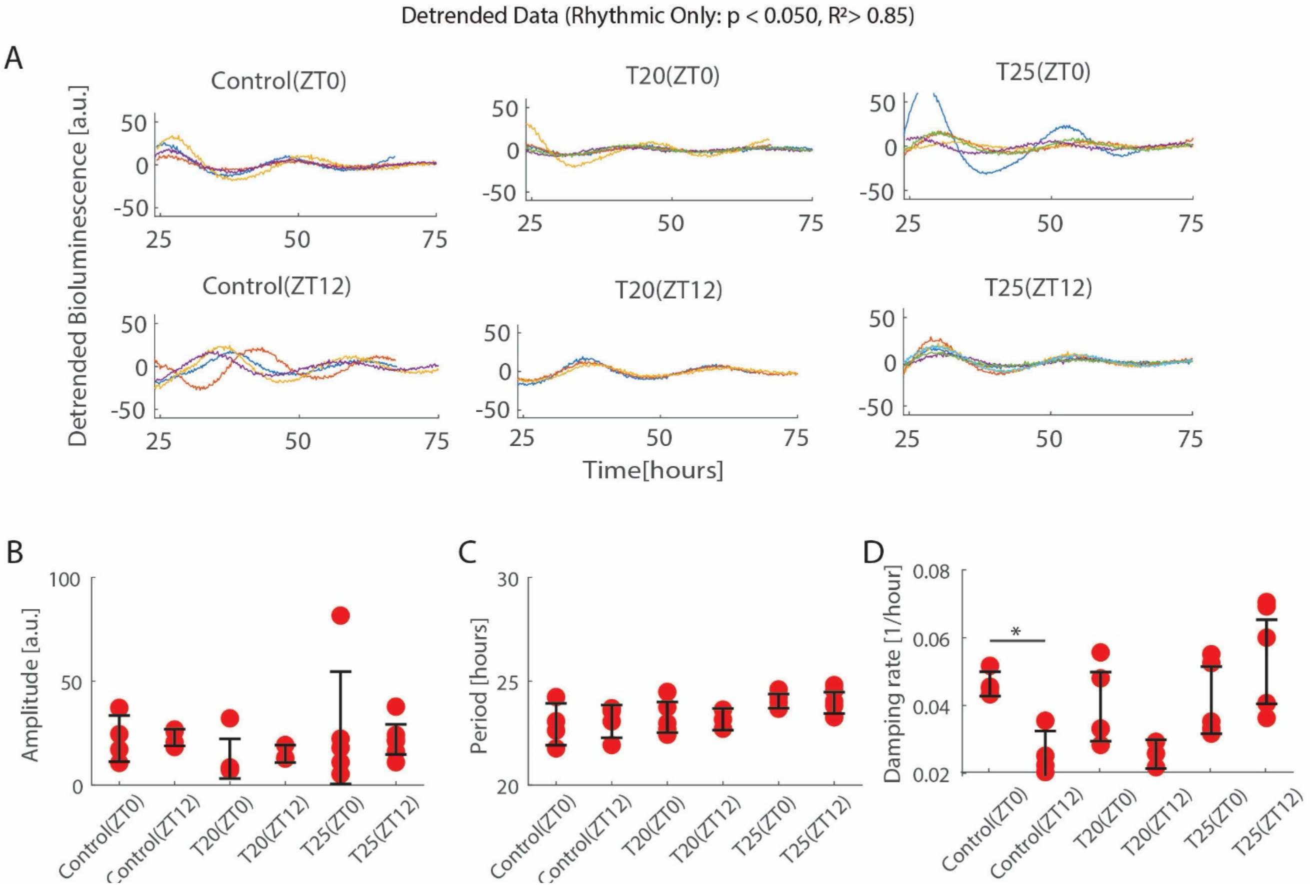


**Supplementary Figure 4. Individual detrended bioluminescence recordings and additional circadian parameters of wounded Dbp:Luc skin explants across entrainment and wounding-times.** A) Detrended bioluminescence recordings (under 37°C) of individual rhythmic wounded Dbp:Luc skin explants for each condition (Control(ZT0), Control(ZT12), T20(ZT0), T20(ZT12), T25(ZT0), T25(ZT12)) from hour 24 to hour 75 after wounding (N≥3). *Time is time since end of T25 entrainment/ZT0 wound. B) Amplitudes, C) periods, and D) damping rates of rhythmic Dbp:Luc skin explants wounded at ZT0 or ZT12 following Control (37°C), T20, or T25 temperature cycle entrainment (mean±SEM; student's t-test, *$p<0.05$).
